# New residential perennial floras reflect distinctive decisions of residential developers and new homeowners

**DOI:** 10.1101/716175

**Authors:** Tracy L. Fuentes

## Abstract

Urban patterns reflect the people who build and manage urban space. However, most field research on residential vegetation focuses on household or neighborhood preferences, norms, or socioeconomic drivers of observed patterns and plant traits. Very few urban ecology researchers have studied residential real estate developers, who configure the space and establish the initial plant communities. How do the landscaping decisions of developers and homeowners shape residential perennial floras? To answer this question, I collected a stratified random sample of perennials at 60 newly built and sold homes in the Seattle, WA area. Through field sampling, conversations with new homeowners, and archival research, I assigned each individual perennial to one of three origins: remnant, planted by developers, or planted by homeowners. After describing landscaping decisions using plant traits (as presented in gardening literature), I evaluated whether planting decisions of developers and homeowners were heterogeneous and whether urban form or economic drivers influenced planted species richness. I also tested whether homeowner yard and plant buying preferences could be linked to planted richness. Given that developers and homeowners have different incentives, I hypothesized that they would choose different types of perennials and that urban metrics related to area and economics would increase species richness. I also predicted that homeowner preferences would be linked to species richness patterns. Developers planted most of the trees, shrubs, and graminoids. Homeowners planted fewer woody and more herbaceous perennial species. Parcel planting area, wealth related metrics, and parcel density increased species richness for some perennials. However, homeowner preferences were stronger predictors of their planting behavior than urban metrics. Because assembly of residential perennial flora communities is heterogeneous, future investigations in other urban ecosystems should incorporate preferences of developers and homeowners, site-specific constraints, and broader scale influences. More work is needed to understand developer incentives and preferences.

## Introduction

Most urban vegetation consists of privately managed residential yards and gardens [1-3]. Given the large land area occupied by single-family residences, understanding residential vegetation dynamics in general requires a careful assessment of how yards and gardens are built and maintained. As Li et al. [4] noted, residential landscapes “say something about the people who construct and use them.” Therefore, examining their structure and composition requires consideration of the human and the biophysical drivers [5-6] operating and interacting at multiple scales [7-9].

Urban metrics linked to urban plant diversity and abundance patterns include neighborhood wealth [10-12], housing age [10,13], and density [14-18]. Wealth-related metrics, in particular, have been linked with increased plant richness and abundance, while housing characteristics and density-related metrics seem to be more varied, depending on the urban system context. The “luxury effect” [10] or “ecology of prestige” [19], where wealthier people have access to more diverse and more abundant vegetation appears to result from complex interactions of individuals, land developers, land owners, institutions, and feedbacks between them [20-22]. Neighborhood wealth has been linked to increased plant species richness [22,23], tree species richness [11], cover and remotely sensed greenness [12, 24].

In addition to wealth metrics, other household demographic factors are associated with landscape preferences and yard management. Cultural background [25], life stage and gender [26,27], and length and type of land tenure [28,29] have all been shown to influence yard preferences and gardening activities. Larson et al. [30] related these household preferences to the desire for managing these semi-private spaces as “landscapes of leisure”. Thus, yards are designed and managed to meet household needs and preferences: spaces in which to garden, play, entertain family and friends, or relax.

To link these diverse and somewhat ambiguously defined resident preferences to specific plant outcomes, researchers have incorporated plant traits into their studies. The use of plant traits in urban residential landscape studies draws on community ecology, where specific traits are analyzed to see how plant taxa identity and abundance vary in response to biophysical patterns and processes [31,32]. Noting the difficulty in linking common plant ecological traits to how people actually choose plants, Pataki et al. [33] proposed to use “ecosystem traits” to detect what benefits people obtain from plants. Some of the ecosystem service traits linked to human preferences are plant size [34], showy flowers [34-36], edible fruit [33], leaf width [35], and shade [28,37]. However, Conway [38] found that people selected trees for aesthetics or low maintenance reasons, not for provisioning ecosystem services.

I build on this body of work by explicitly evaluating the different plant choices by developers and homeowners in newly built and sold residential landscapes. From an urban ecology perspective, residential vegetation can be seen as the outcome of human decisions interacting with multi-scalar processes [7-9,39,40]. Specifically, two diverse human agents, development firms and homebuyers, interact in the regional real estate market to meet their specific needs and goals.

The initial plant structure and composition of the new landscape reflect this process. Developers compete for and buy land, complete neighborhood and site planning processes, and obtain subdivision and building permits from the local planning authority [41]. The developers then install or upgrade infrastructure, build homes, and add other paved surfaces in and adjacent to the parcel. The landscaping is typically one of the last steps, along with other presale work [42]. Depending on the location, municipalities and/or homeowners associations may specifically regulate aesthetics, planting plans, tree removal, and maintenance for subdivisions and for yards [43-46]. After the new homeowners move in, they may or not update or change the initial yard configuration, remove plants, and plant more or different species.

However, urban ecologists have devoted little attention has been devoted to the real estate developer industry, despite the fact that “it is a highly regulated, high value industry that shapes the built environment” [47]. Thus, most of the literature about residential plant communities focuses on households or neighborhoods [8], despite the fact that developers set the amount and configuration of growing space [42,48,49] and limit which taxa and which life forms can establish after the initial plant community is established [50].

To understand how the complex interactions of developer and homeowner decisions shape new residential floras, I focus specifically on new residential landscapes in south King County, WA. Through site visits, information from homeowners, and archival photo research, I document specific landscaping decisions in the new yards. How do developer and homeowner landscaping decisions shape the new yards? Do they plant the same types of perennials? Do they select the same or different taxa? How diverse are their selections? Are some plant species or families planted more frequently than others? Which urban metrics or household preferences influence planted species diversity?

Although many researchers are attempting to link plant traits to resident choices, none of these specifically consider plant traits that are commonly recommended for choosing plants. Therefore, I reviewed a number of ways that landscape design and gardening books classify all perennials when recommending perennials. I chose Stoecklein’s [51] approach: evergreen and deciduous trees, evergreen and deciduous shrubs, groundcovers, vines, ferns, graminoids, and other herbaceous perennials. Not only does this classification align with urban ecology, plant trait, and landscape design literature, it also is consistent with how plants are displayed and marked in stores and nurseries. For example, mature height and leaf habit (deciduous or evergreen) have been found to affect tree community structure and composition in residential landscapes and are mentioned as considerations in tree preferences [28,36,37,52,53], landscape design [51,54] and gardening books [55,56].

First, I describe and compare the remnant, developer selected, and homeowner selected perennials. How large and how diverse are these species pools? What proportion is native to the Pacific Northwest? I predict that there will be structural and compositional differences in these pools. The two agents will select different types and taxa, but both will favor certain structural groups or taxa more than others.

Second, for both developers and homeowners, I test to see if there are specific urban metrics linked to the diversity patterns grouped by plant buying structural groups. I predict that metrics at the site and neighborhood scale will be associated with plant structural group richness. In particular, wealth-related metrics should be positively associated with increased diversity. At the site scale, these are appraised improvement value and land value. At the neighborhood scale, this is median assessed value, which includes both land and building values.

Third, I test whether expressed homeowner plant selection criteria or yard management goals are reflected in specific patterns in the homeowner planted perennials. I hypothesize that yard management goals and plant selection favoring easier to manage landscapes and species should be negatively associated with woody perennials, while those relating to providing food and wildlife habitat should be positively associated with richness. Other yard management and selection criteria may also be influential but may vary by group.

Fourth, I compare the composition and diversity of the presale and postsale plant communities. I define the presale community as resulting from developer decisions to keep specific plants on the parcel and to plant new perennials after the house and other infrastructure is completed. The postsale community consists of plants from the presale flora that the homeowner kept, as well as new perennials that they plant. I predict that taxa diversity will increase in the postsale flora, but not for all groups.

By focusing solely on perennial plants in new yards of new homes, I identified the specific decisions of each agent and described the initial floristic conditions of these new floras. I found broad support for my hypotheses that developers and homeowners select different plant structural groups and species and that their choices reveal different preferences. Developers plant more woody perennials and graminoids, and homeowners plant more herbaceous perennials. Thus, the majority of woody and graminoids perennial abundance and richness in new yards in this sample results from decisions of developers, while the rest of the herbaceous flora more strongly reflects homeowner decisions.

## METHODS

### Regional context

The Puget Sound Region of Washington State supports diverse terrestrial communities resulting from the abrupt elevation, temperature, and moisture gradients [57]. Lowland vegetation (<150m) on the west slope of the Cascades Mountains is mostly temperate coniferous forest. Dominant conifers are *Pseudotsuga menziesii* (Douglas-fir), *Tsuga heterophylla* (western hemlock), and *Thuja plicata* (western red-cedar). Less common are *Quercus garryana (*Garry oak) woodlands and prairies in the southern part of the region, which have been greatly reduced from their historical extent from agriculture and then urbanization.

The Seattle-Tacoma-Bellevue-Metropolitan Statistical Area (Seattle MSA) occurs in the central part of the Puget Sound Region and contains all of King, Pierce, Snohomish, and Kitsap Counties [58]. An estimated 3.87 million people live in the Seattle MSA, with the majority (2.2 million) living in King County.

### Sampling frame, sample, and recruitment

The sampling frame (N=1,258) is newly built and sold single-family residential parcels within the lower Green-Duwamish watershed in King County, WA, USA (Fig 1). I chose this study area for several reasons: 1) it contains the Duwamish River, Seattle’s only river; 2) the heavily industrialized watershed contains the Port of Seattle, SeaTac airport, and three of eight designated Manufacturing and Industrial Centers in the Puget Sound Area [59]; 3) parts of the watershed contain diverse, low-income neighborhoods [60]; and 4) high quality parcel data and real estate transactions are available [61].

**Fig 1.**
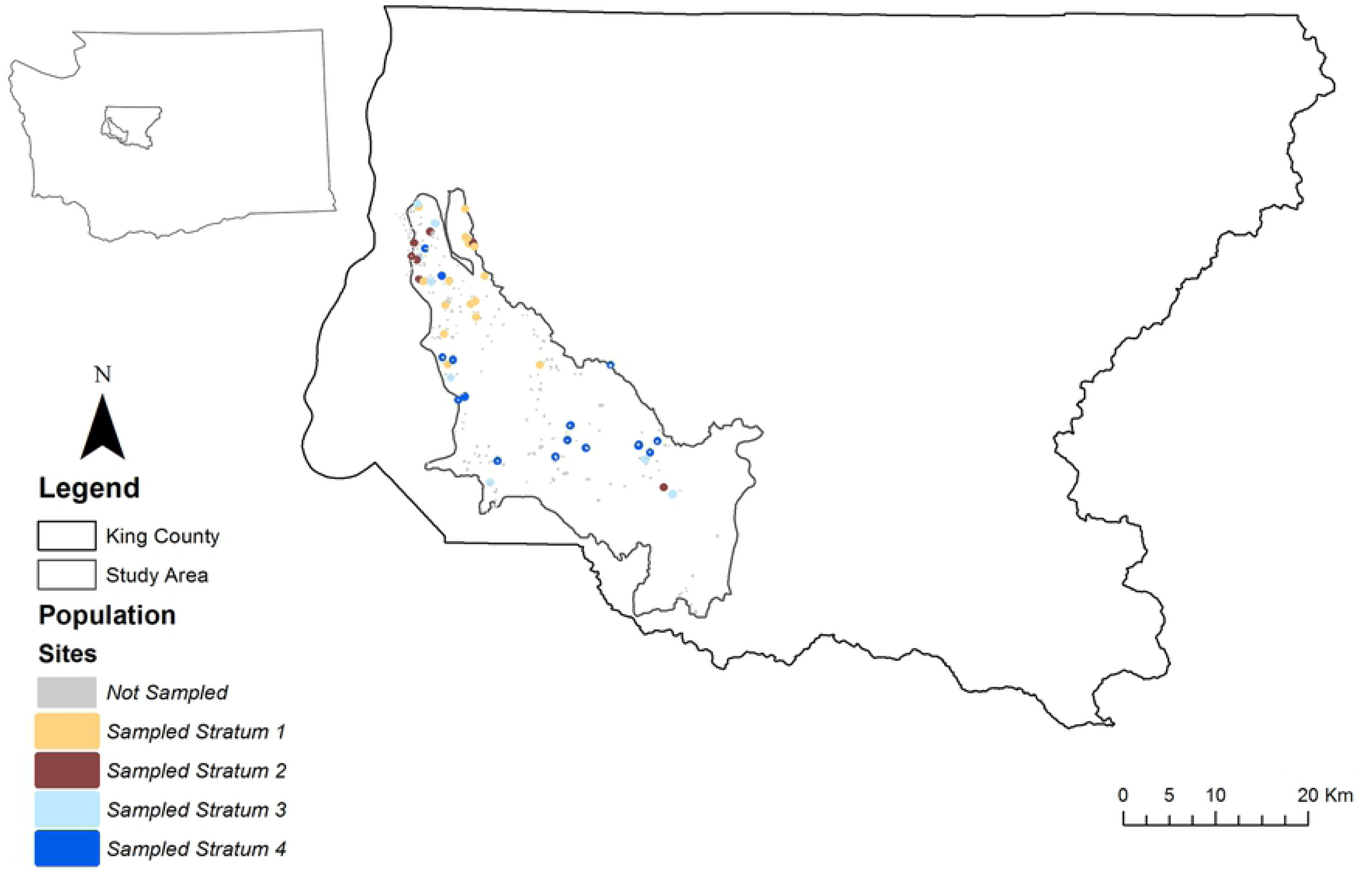
Location of the lower Green-Duwamish study area and sampling frame parcels within a) Washington State and b) King County. Parcels in grey are part of the sampling frame but were not sampled. Sampled parcels are from one of four strata: 1= developer has 1 parcel in population; 2 = developer has 2 parcels in population; 3= developer has 3-8 parcels in population; 4 = developer has 9 or more parcels in population.

To be eligible for inclusion in the sampling frame, parcels had to meet all of the following criteria: 1) be located within the study area, 2) have a single family home built in 2014 or 2015, 3) be sold by a developer to a household by July 1, 2016. By requiring that the owner be an individual or a household, I eliminated parcels that are held by firms. By requiring that the builder be different than the household, I eliminated households who have built their dream homes and yards.

The study sample is a stratified random sample of 60 parcels in the sampling frame (Table 1). Strata are derived from quartiles of parcel counts for each unique development firm. Strata 1 and 2 represent infill development and small developers, who have built and sold 1 or 2 parcels, respectively, in the sampling frame. Strata 3 and 4 represent small subdivision developers (3-8 parcels) and large subdivision developers (9+ parcels). Within King County [62], a short plat subdivides property into four or fewer parcels outside the Urban Growth Area and nine or fewer parcels within the Urban Growth Area. Thus, natural breaks in the dataset follow actual building practice.

**Table 1.** Responses by stratum and recruitment method. Mail was random sample of 40 parcels/stratum. In Person #1 is in-person visit to parcels previously mailed. In Person #2 is in-person visit to new draw of 20 parcels/stratum. However, I did not visit all 20, because I stopped when I reached 15 affirmative responses per stratum. Yes = agreed to participate; No = refused to participate; NR = no response by mail or unable to contact homeowner in person.

| Population |  |  | Responses by Recruitment |  |  |
| --- | --- | --- | --- | --- | --- |
| Stratum | #Parcels/<br>Developer | # Parcels | Mail<br>Yes/No/NR | In Person #1<br>Yes/No/NR | In Person #2<br>Yes/No/NR |
| 1 | 1 | 126 | 9/1/30 | 4/3/23 | 2/4/2 |
| 2 | 2 | 56 | 7/2/31 | 5/2/24 | 3/2/4 |
| 3 | 3-8 | 131 | 10/6/24 | 1/1/22 | 4/0/11 |
| 4 | 9+ | 945 | 10/2/28 | 2/0/26 | 3/8/6 |

To obtain permission to sample, I sent a letter to the registered owner describing the research. To increase likelihood of response, I personally signed each letter, hand wrote the address, and included a self-addressed stamped post-card [63]. Because of low response rates (Table 1), I dropped my target number of parcels per stratum from 25 to 15. To fill the remaining slots per stratum, I visited the potential sites to recruit in person, on a Saturday or Sunday, between 10am-5pm. I first visited the homes that I previously mailed. Then, I selected a new random sample of 20 parcels from each stratum, after attempting to contact from first random draw. I stopped recruiting from random sample two when I reached a full set of 15/stratum.

### Documenting perennials and source

During the growing season of 2016, I conducted a complete vegetation survey of all perennial plants within the actively managed area of the sample parcels. I defined this area as the buildable portion of the parcel, from the street to identifiable property boundaries, such as fences or sidewalks and driveways of adjacent parcels. To conduct the survey, I assigned each planting bed a unique patch number, then identified and counted all perennials within that patch.

I assigned each individual or clump of individual plants as belonging to one of three origin categories: remnant, developer planted, or household planted. Remnant perennials are those that were on site predevelopment and were retained by the developer. They typically include large trees, shrubs, and some rock wall species. Developer planted individuals are 1) planted perennials that the homeowners said were present when they bought the house, 2) planted individuals that the homeowners told me they removed, 3) planted individuals that were present in images prior to sale date and that were absent during the site visit. Household planted individuals are those that the household informed me they planted. Large native trees would be classified as remnant, but their seedlings would be classified as spontaneously establishing on site and were excluded.

Because my research questions focus on developer and homeowner landscaping decisions at the parcel level, I used patches, not plots, as the sampling unit. For standard geometric shapes, I measured patch dimensions necessary for area calculations. For irregular patches, I used the offset method [64]. To correct for any measurement error and differences in parcel area versus sampled area, I corrected yard area and total area of each patch type via aerial photos analysis of the sampled parcels. Using measuring tools and aerial photos available at the King County Parcel Viewer [65], I measured areas of the building footprint, impervious surfaces, and the remainder of the parcel that was developed. This excludes steep slopes and wetlands and irregular areas outside of fences but includes curb strips. Corrected yard area is the difference of sampled area and all impervious surfaces within it.

At the time of the field survey and afterwards, I asked the new homeowners if they had added or removed plants. To obtain additional information about what plants may have been present earlier, I compiled images of the parcel before the sale [65, 66, 67]. Nomenclature follows Hitchcock et al. [68] or Missouri Botanic Garden [69].

### Homeowner preferences

People select plants based on they how they want to use the space and specific characteristics of the plants themselves [29, 33, 38, 52, 56, 57]. Therefore, I mailed participating households a yard management questionnaire after the site visit. Of the 60 homes in the original sample, 35 responded to my yard management questionnaire. Because one of these is a parcel I discarded from further analysis in this paper, final sample size for household preference analyses is 34.

Among other questions, I asked each household to rate the importance of 10 plant selection criteria and eight different yard management goals. Rated yard management goals included variables to incorporate aesthetics (*Beautiful, Neat*), ease of maintenance (*Easy*), food production (*FoodSpace*), provision of wildlife habitat (*Habitat*), and other uses of space (*Entertain, KidSpace, PetSpace*). Rated plant selection criteria included variables to measure subjective criteria (*Beauty, Unique, Personal*), price criteria (*Inexpensive*), food and wildlife related criteria (*Household Food, Nectar, Wildlife*), as well as maintenance criteria (*Low Maintenance)* and drought tolerance (*Drought*).

### Data management and analysis

After I entered the field and questionnaire data, I rechecked all values a second time. All further data manipulation conducted in R 3.5.1 [70] and *tidyverse* [71].

After final plant identifications, I placed each taxon within one of nine perennial structure groups, following Stoecklein [51]: evergreen or deciduous trees, evergreen or deciduous shrubs, graminoids, groundcovers, ferns, vines, and all other herbaceous perennials (S1).

Trees are tall woody perennials that are >2m at mature height. I included *Trachycarpus fortunei* (windmill palm) as a tree, rather than other perennial monocot, because that is how it is recommended to landscapers and gardeners in the Pacific Northwest. Shrubs include woody taxa <2m in height. Graminoids are true Poaceae, as well as Cyperaceae, Juncaceae, Typhaceae, Asphodelaceae, and a subset of Asparagaceae that are recommended as grasses. Groundcovers are low growing perennials recommended for covering soil and preventing weed establishment. Ferns are members of the Dryopteridaceae. Vines have flexible stems that can be trained around supports. The other perennial group includes all remaining dicot and monocot taxa. These include perennial flowers, small herbs, and unusual genera that don’t easily fit into the other categories (e.g. *Cynara, Sedum, Musa, and Cycas*).

For all parcels, I compiled and calculated variables representing urban form, gradients, and land economics (Table 2). Parcel variables are from on-site measurements or from King County [61]. Neighborhood variables are at the enclosing Census Block Group extent (hereafter CBG), which is commonly used in urban vegetation studies [12,37,72]. However, I calculated all estimates directly from parcel data [61] for each CBG that contained a sampled yard.

**Table 2.**
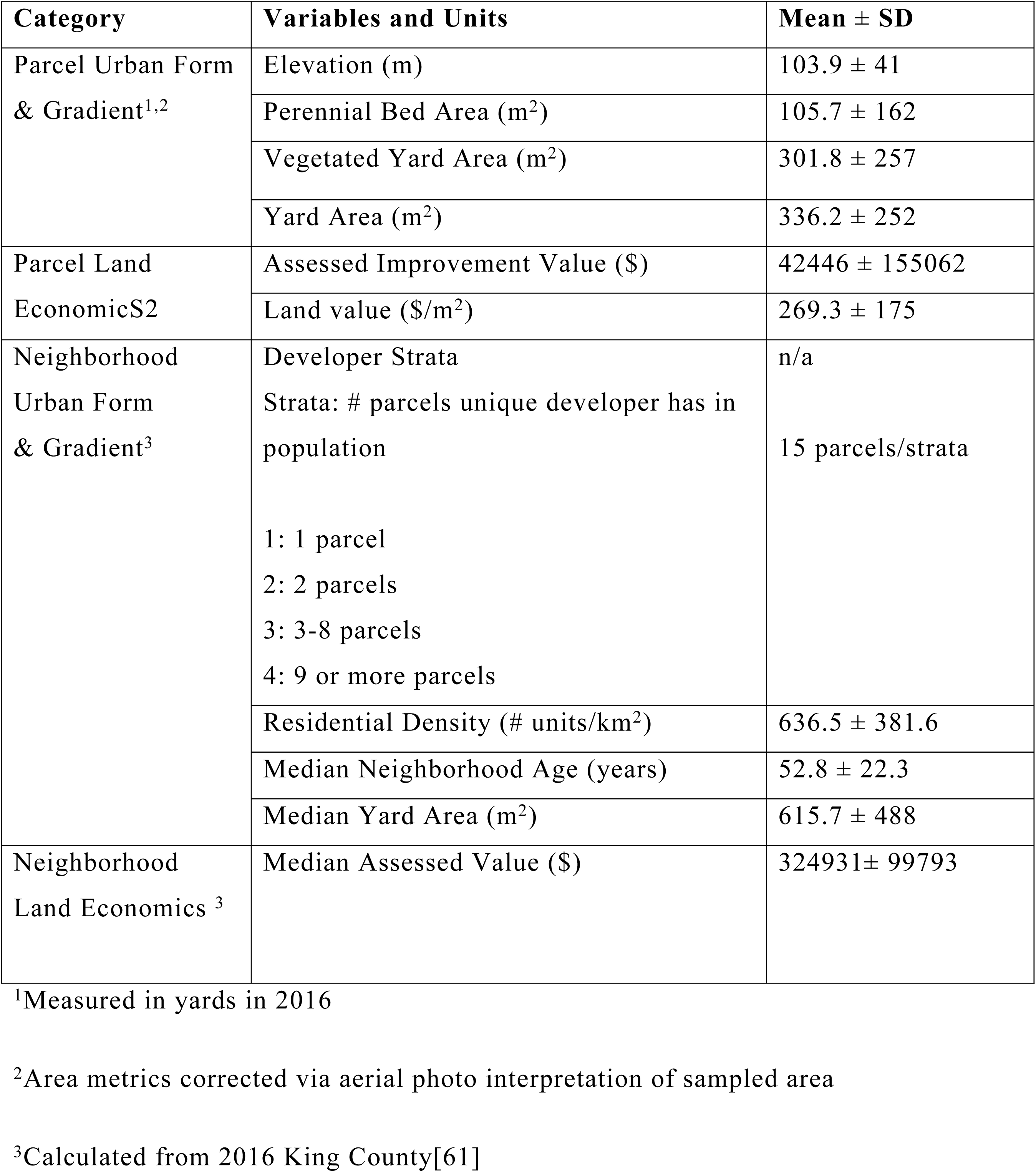
Independent variables in negative binomial regressions of planted species richness, by category, name and characteristics (n=58).

To calculate diversity indices, I used *vegan* [73]. I used *MASS* [74] for multivariate regressions. I tested for spatial autocorrelation via Monte Carlo simulation of Moran’s I in *spdep* [75] and fit models with spatial lags when p<0.1. Numeric independent variables in the parcel, neighborhood, and household models are mean centered and divided by the standard deviation. Final models are those with the smallest corrected Akaike (AIC_c_) values, which were calculated in *AICcmodavg* [76]. To calculate model p values, I conducted likelihood tests of potential final models by comparing them to next simpler models.

To test evaluate planting decisions by agent, between agents, or with planting area, I calculated Spearman rank correlations in *Hmisc* [77] and plotted them with *corrplot* [78]. Then, I ran multilevel logistic regressions using *lme4* [79] and *mlmRev* [80]. Fixed effect is plant structural group. Random effect is the agent interacting with the site, to control for area and other unique aspects of the parcel. Each unique taxon present on the parcel was coded as 1 for that group. Absent structural groups were coded as 0.

To assess which urban metrics and household preferences influenced planted species richness, I fit full negative binomial regressions and allowed stepwise regression (both directions) to suggest the best fitting model. I evaluated model fit using corrected AIC and likelihood tests as above. For each agent, I fit a full site model and a full neighborhood model for tree, shrubs, and herbs. Then, I fit full household yard and plant selection criteria models. Because of the small sample size and correlations in preference variables, I used a reduced set of preferences as independent variables (Table 3).

**Table 3.**
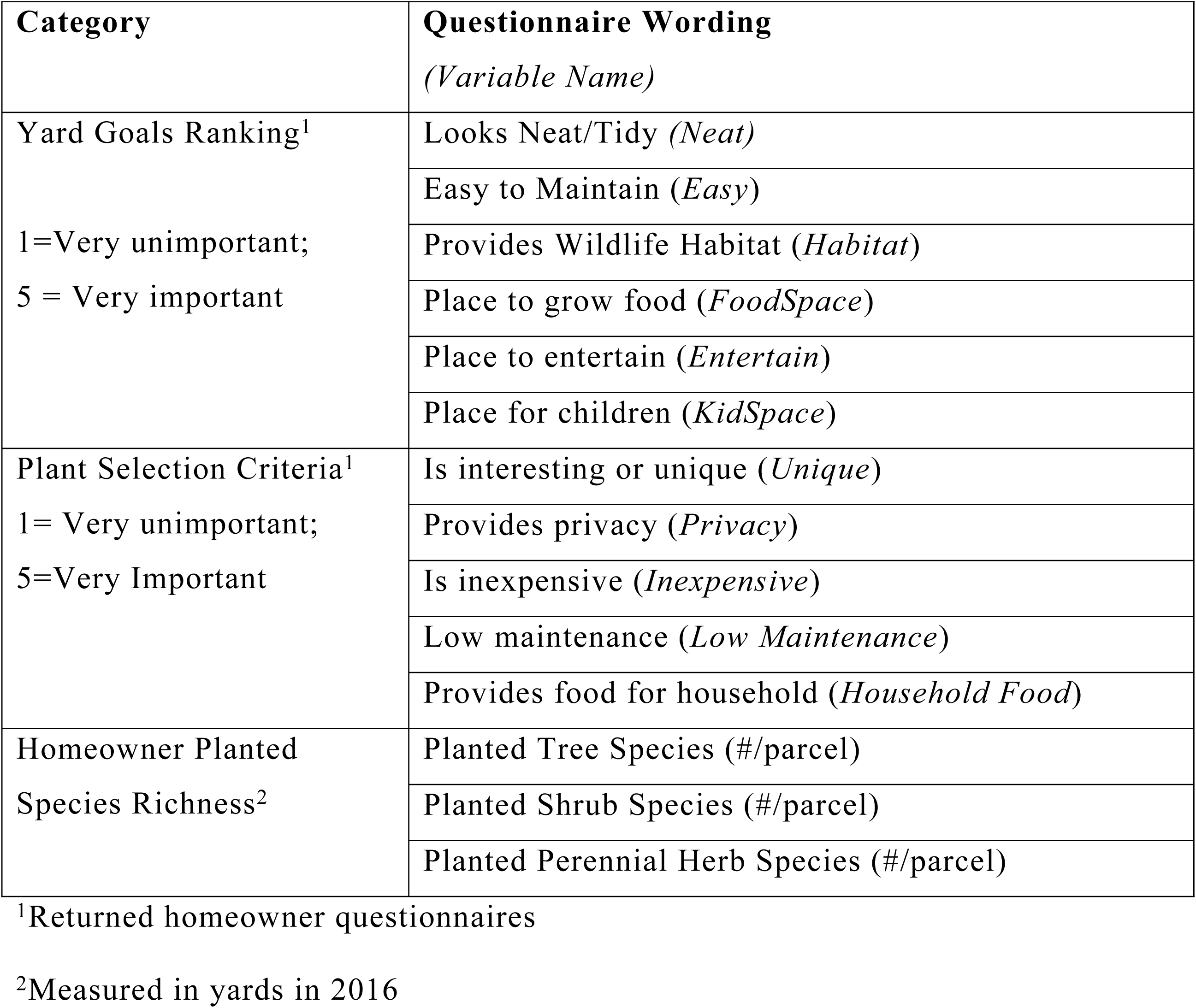
Independent variables used in negative binomial regressions of homeowner planted species richness (n=34).

| Category | Questionnaire Wording<br>(Variable Name) |
| --- | --- |
| Yard Goals Ranking<sup>1</sup><br>1=Very unimportant;<br>5 = Very important | Looks Neat/Tidy (*Neat*) |
|  | Easy to Maintain (*Easy*) |
|  | Provides Wildlife Habitat (*Habitat*) |
|  | Place to grow food (*FoodSpace*) |
|  | Place to entertain (*Entertain*) |
|  | Place for children (*KidSpace*) |
| Plant Selection Criteria<sup>1</sup><br>1= Very unimportant;<br>5=Very Important | Is interesting or unique (*Unique*) |
|  | Provides privacy (*Privacy*) |
|  | Is inexpensive (*Inexpensive*) |
|  | Low maintenance (*Low Maintenance*) |
|  | Provides food for household (*Household Food*) |
| Homeowner Planted<br>Species Richness<sup>2</sup> | Planted Tree Species (#/parcel) |
|  | Planted Shrub Species (#/parcel) |
|  | Planted Perennial Herb Species (#/parcel) |

<sup>1</sup>Returned homeowner questionnaires
<sup>2</sup>Measured in yards in 2016

To assess how composition and flora diversity changes by ownership, I created two floras. Presale flora consists of perennials present at the time of sale. Postsale flora consists of plants present at the time of site visit and excludes removed or dead perennials. All diversity indices are in effective species number equivalents [81,82]: richness, exponent (H’) and 1/Simpson. By converting them to taxa equivalents, the indices exhibit common behaviors allowing for comparing indices in an ecologically meaningful manner. To properly partition taxonomic diversity, I used the multiplicative versions of gamma, alpha, and beta diversities [83,84]. Gamma diversity is the total count of unique species, genera, or families. Alpha and beta diversity are calculated twice: by mean taxa richness and mean exponent (H’). Beta is gamma divided by alpha.

## RESULTS

### Remnant taxa

Total number of remnant taxa is 38 species, 33 genera, and 20 families (Table 4, S1). Of these remnant taxa, 39% were native to the Pacific Northwest, and 61% were not. The six most frequent remnant families are Pinaceae, Rosaceae, Ericaceae, Cupressaceae, Dryopteridaceae, and Sapindaceae (pine, rose, heather, cedar, wood fern, and soapberry families, respectively. Three of the most frequent species were large remnant trees (*Pseudotsuga menziesii* (Douglas-fir), *Arbutus menziesii* (Pacific madrone), and *Prunus serrulata* (Japanese flowering cherry). The native *Polysthichum munitum* (sword fern) was the other frequent remnant species. All other taxa occurred at two or fewer sites.

**Table 4.** Comparison of perennial sources in new residential landscapes by number of individuals, species, genera, and plant families. Six most frequent families are arranged from most to least frequent.

| Source | Individuals | Species<br>(# native) | Genera | Families | Six Most Frequent<br>Families |
| --- | --- | --- | --- | --- | --- |
| Remnant | 174 | 38<br>(15) | 33 | 20 | Pinaceae, Rosaceae,<br>Ericaceae,<br>Cupressaceae,<br>Dryopteridaceae,<br>Sapindaceae |
| Developers | 2483 | 108<br>(23) | 78 | 41 | Cupressaceae,<br>Ericaceae, Cyperaceae,<br>Sapindaceae, Poaceae,<br>Berberidaceae |
| Homeowners | 905 | 134<br>(14) | 106 | 60 | Rosaceae,<br>Asparagaceae,<br>Ericaceae, Iridaceae,<br>Lamiaceae,<br>Cupressaceae |

### Planting behavior

Developers planted most of the perennials on each parcel (Table 4; S1). About 21% of the developer pool (101 species) is native to the Pacific Northwest, compared to 10% of the homeowner pool (127 species). Of the 58 homeowners, 12 planted no perennials.

Developers and homeowners showed distinct differences for structure groups when selecting new species to plant (Fig 2, 3). Developers had much greater mean probabilities (>0.88) of planting tree, shrub, and graminoid species than homeowners (mean probability 0.13-0.27). Of the woody species, developers favor evergreen species over deciduous.

**Figure 2.**
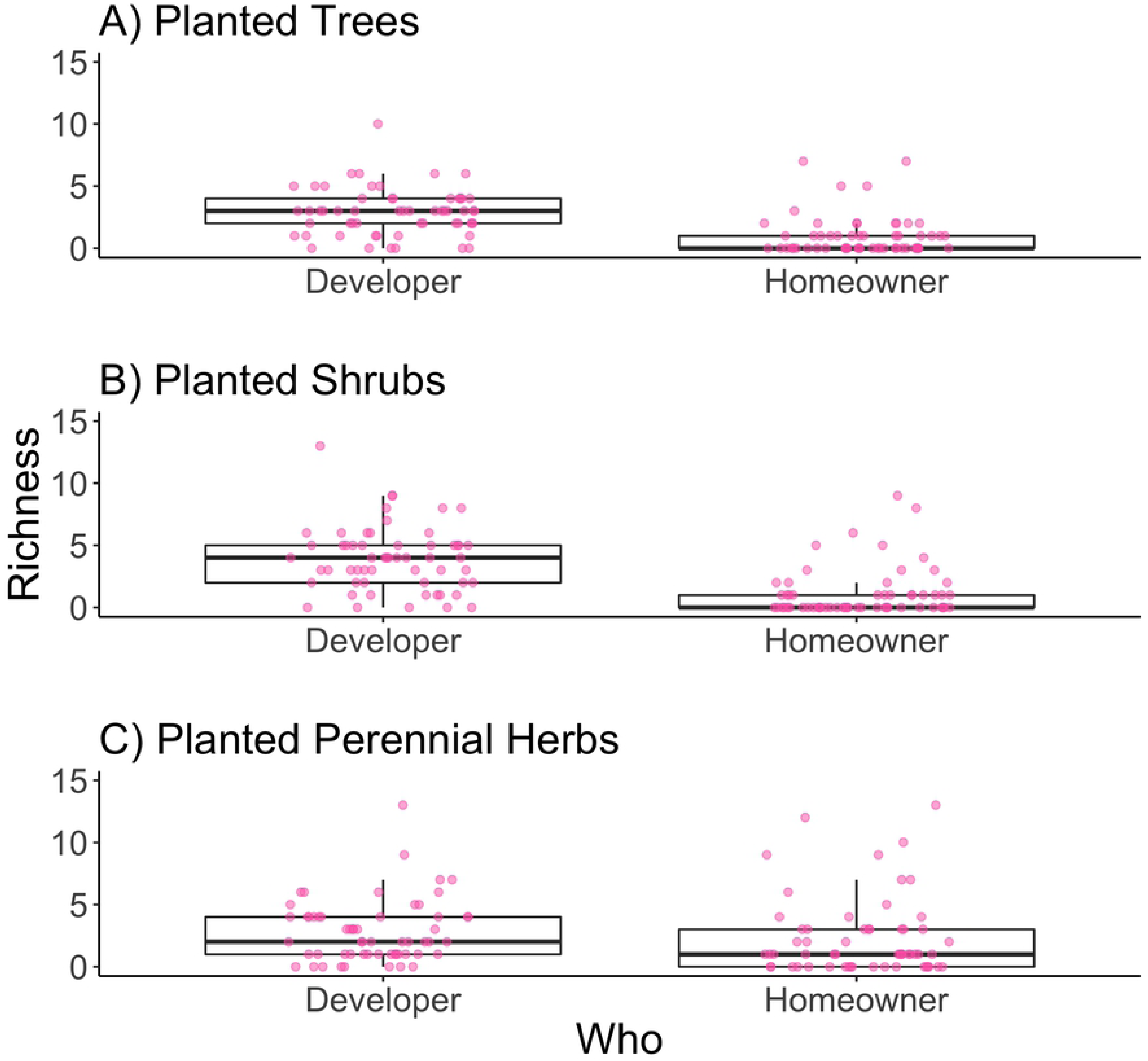
Boxplots of developer and homeowner planted richness for A) trees, B) shrubs, and C) perennial herbs. Points are species richness at each site, jittered horizontally to spread similar values.

**Figure 3.**
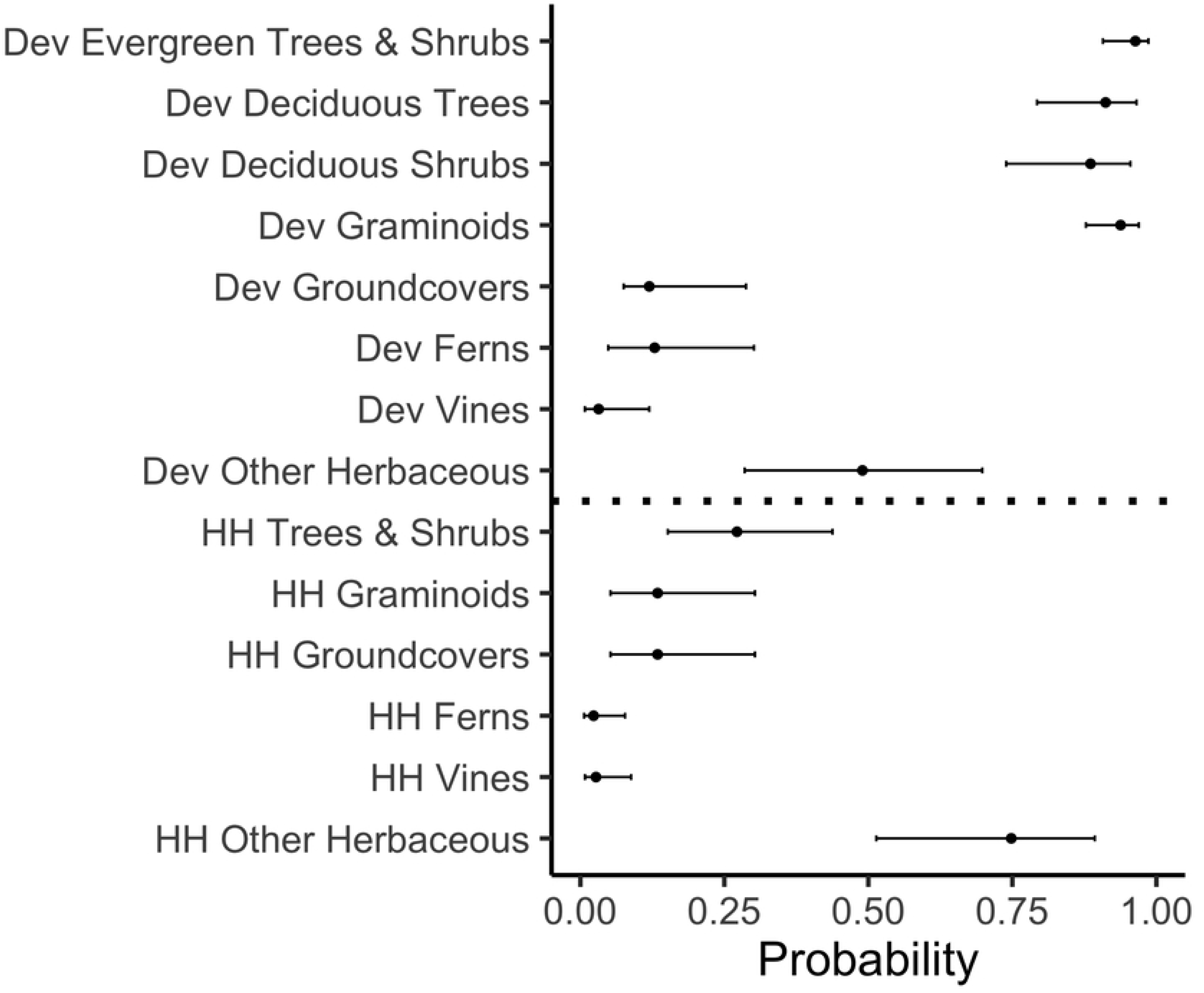
Probabilities that developers and homeowners chose specific plant structure groups (Stoecklein 2011). Differences in area and other unique characteristics of the yard and location controlled as a random effect in the mulitlevel models (S2). Closed circles represent the mean probability that the developer or homeowner selected that group. Error bars are calculated 95% confidence intervals. Developer plants begin with “Dev” and are above the dotted line. Homeowner plants begin with “HH” and are below the dotted line. Developer evergreen trees and shrubs are combined, as are homeowner trees and shrubs; models detected no difference in probability for those groups).

Homeowners showed no difference for evergreen habit or for trees or shrubs when choosing to plant a woody species. Instead, homeowners select “other herbaceous” more than any other category. Homeowners had a greater mean probability (0.75) of selecting species in the “other herbaceous category than developers did (0.49).

### Urban metrics of species richness

Planting behavior of each agent is a strong correlate of richness, as are various metrics of planting area (Fig 4). Developer tree richness is positively correlated with developer planted shrub (Spearman rho=0.45) and herbaceous perennial richness (Spearman rho =0.25). Similarly, homeowner tree richness is positively correlated with homeowner shrub (Spearman rho=0.44) and herbaceous perennial richness (Spearman rho =0.44). Developer and homeowner planted richness are not correlated.

**Figure 4.**
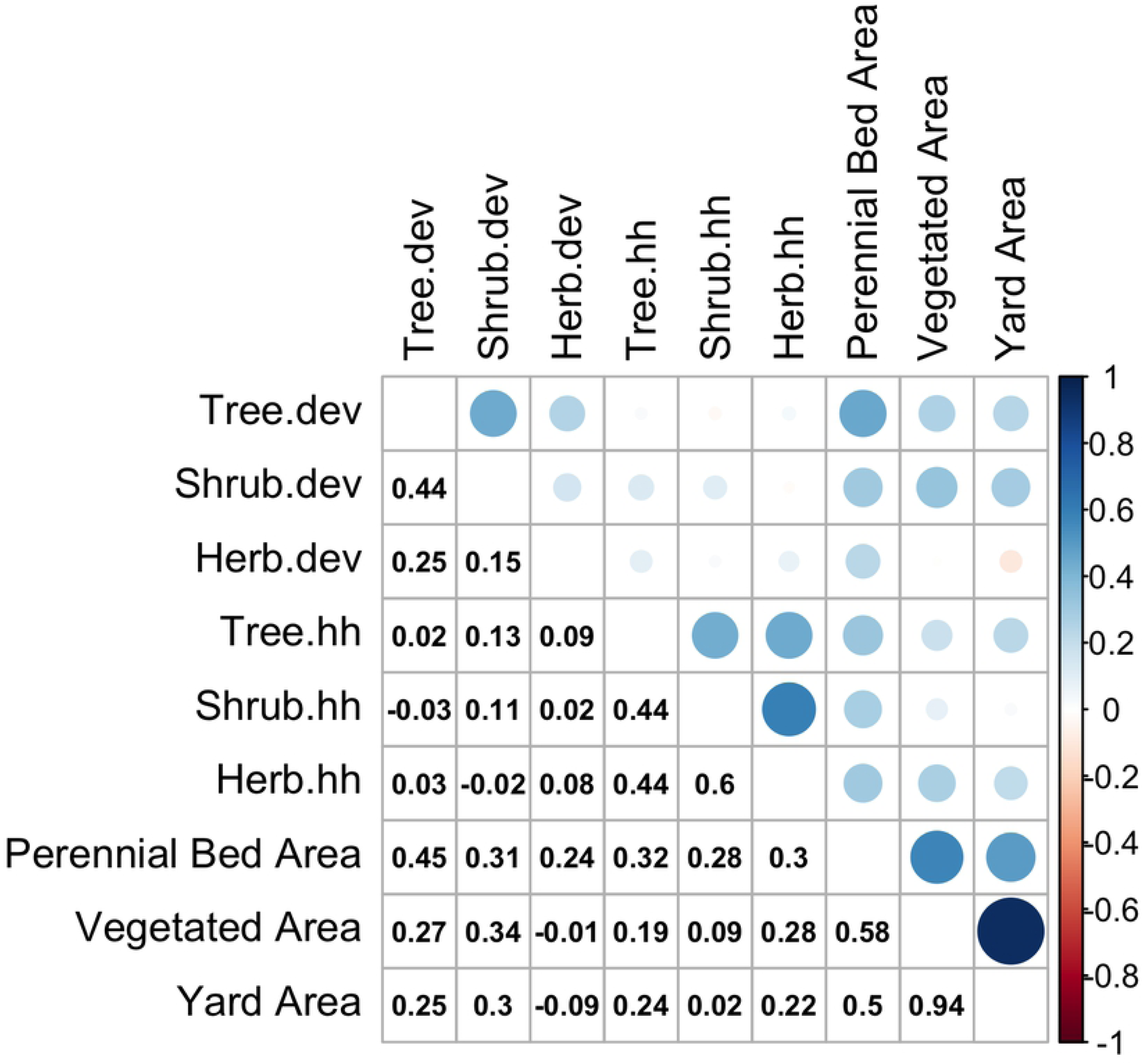
Spearman rank correlations between planted perennial richness and area. Developer planted species richness for trees, shrubs, and herbs ends in “.dev”; homeowner planted richness ends in “.hh”. Perennial Bed Area = total area of all perennial beds (m^2^); Vegetated Area = sum of perennial bed area and lawn area (m^2^); Yard Area = pervious area of developable parcel area (m^2^). Size of circles indicates strength of correlation. Blue circles indicate positive correlations; red circles indicate negative correlations.

**Figure 5.**
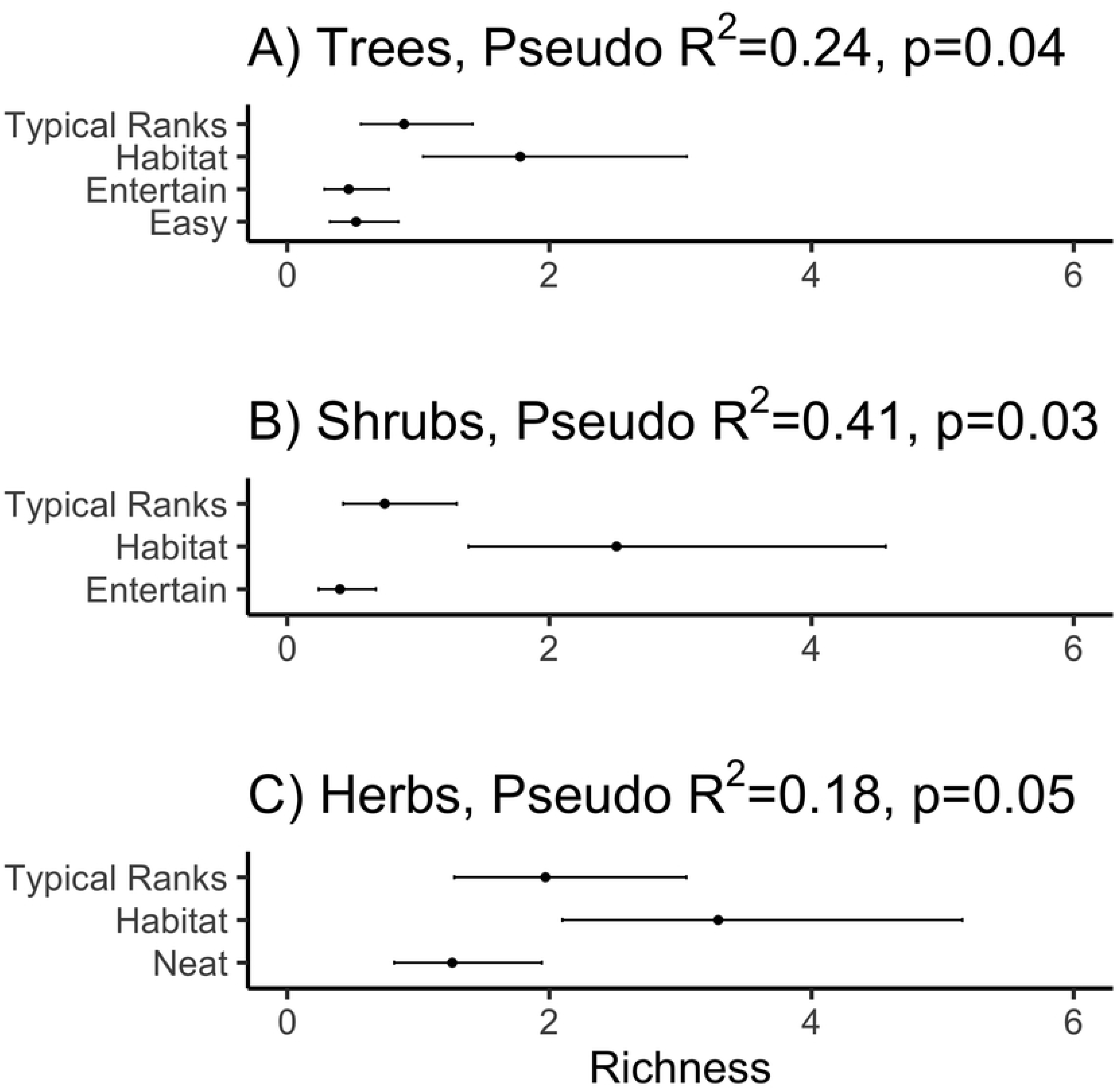
Estimated effects of higher rankings of yard goals on developer planting behavior for A) trees, B) shrubs, and C) perennial herbs (n=34). Richness is the number of species that the homeowner planted. Closed circles represent the mean species richness of homeowner planted perennials from final regression models (Table 6; S3). Error bars are 95% confidence intervals. *Typical Ranks*: Typical homeowner ranking of goals; *Easy*= Ranking Easy higher than typical; *Habitat* = Ranking Habitat higher than typical; *Neat* = Ranking Neat higher than typical.

Although planted area clearly influenced planted species richness, so did economic drivers and parcel density (Table 5,6). Urban form and economics influenced the planting decisions of developers and homeowners, but these models explained <20% of the deviation. Parcel models explained the most deviation for trees (11-13%), shrubs (7-10%), and homeowner planted herbaceous perennials (6%). For developers, the neighborhood model explained more deviation for developer planted woody perennials (19%). Only area influenced developer shrub richness, homeowner tree richness, and homeowner herb richness

**Table 5.** Standardized coefficients of developer planted species richness from best fitting negative binomial regression models of parcel and neighborhood (Census Block Group) urban metrics (S2). Pseudo R^2^= estimate of deviation explained by model; P = probability from likelihood test compared to next simplest model. AIC_c_ = Akaike Information criterion corrected for small sample sizes). AIC_c_ values in the same column can be compared. Because Monte Carlo simulation of Moran’s I was <0.08 for developer shrubs and >0.5 for all other developer planted perennials, developer shrub models have spatial lags but other models do not.

| **Final Developer Parcel Model** | **Trees** | **Shrubs** | **Herbs** |
| --- | --- | --- | --- |
| Appraised Improvement Value (\$) | 0.153 | | 0.2153 |
| Perennial Bed Total Area (m<sup>2</sup>) | 0.156 |  |  |
| Yard Area (m<sup>2</sup>) |  | 0.1674 |  |
| Spatial Lag |  | 1.0078 |  |
| Pseudo $R^2$ | 0.129 | 0.101 | 0.567 |
| P | 0.043 | 0.054 | 0.053 |
| $AIC_c$ | 231.1 | 269.4 | 250.8 |
| **Final Developer Neighborhood Model** | **Trees** | **Shrubs** | **Herbs** |
| Residential Density |  |  | 0.3756 |
| Median Assessed Value | 0.193 |  |  |
| Spatial Lag |  | -1.1821 |  |
| Pseudo $R^2$ | 0.088 | 0.0503 | 0.06 |
| P | 0.013 | 0.062 | 0.053 |
| $AIC_c$ | 231.9 | 270.8 | 243.3 |

**Table 6.** Standardized coefficients of homeowner planted species richness from best fitting negative binomial regression models of parcel and neighborhood (Census Block Group) urban metrics (S2). Pseudo R^2^= estimate of deviation explained by model; P = probability from likelihood test compared to next simplest model. AIC_c_ = Akaike Information criterion corrected for small sample sizes). AIC_c_ values in the same column can be compared. Because Monte Carlo simulation of Moran’s I >0.60 for all homeowner planted perennials, no models have spatial lags.

| **Final Homeowner Parcel Model** | **Trees** | **Shrubs** | **Herbs** |
| --- | --- | --- | --- |
| Appraised Improvement Value (\$) | | | |
| Perennial Bed Total Area (m<sup>2</sup>) |  | 0.367 |  |
| Vegetated Area (m<sup>2</sup>) |  |  |  |
| Yard Area (m<sup>2</sup>) | 0.416 |  | 0.323 |
| Pseudo $R^2$ | 0.112 | 0.085 | 0.066 |
| P | 0.012 | 0.034 | 0.053 |
| $AIC_c$ | 158.8 | 170.2 | 250.8 |
| **Final Homeowner Neighborhood Model** | **Trees** | **Shrubs** | **Herbs** |
| Residential Density |  |  |  |
| Median Assessed Value |  | 0.3466 |  |
| Pseudo $R^2$ | 0 | | 0 |
| P | 1 | 0.058 | 1 |
| $AIC_c$ | 162.8 | 171.1 | 248.5 |

I detected a luxury effect, where urban economic variables increased richness for some groups. Appraised land value was never selected in models, but appraised improvement value and median assessed value were. Appraised improvement value increased species richness for developer trees and perennial herbs. Median assessed value increased developer tree richness and homeowner shrub richness. However, for perennial herbs, the neighborhood model with parcel density explained more deviation than the parcel model with appraised improvement value.

### Household preferences as drivers of planted perennial richness

Homeowner preferences for how they want to use their yard and choosing plants influenced planting decisions (Table 7; Figs 6,7). Yard goal models explained 18-40% of the deviation in planted species richness, and planting criteria models explained 15-39%.

**Table 7.**
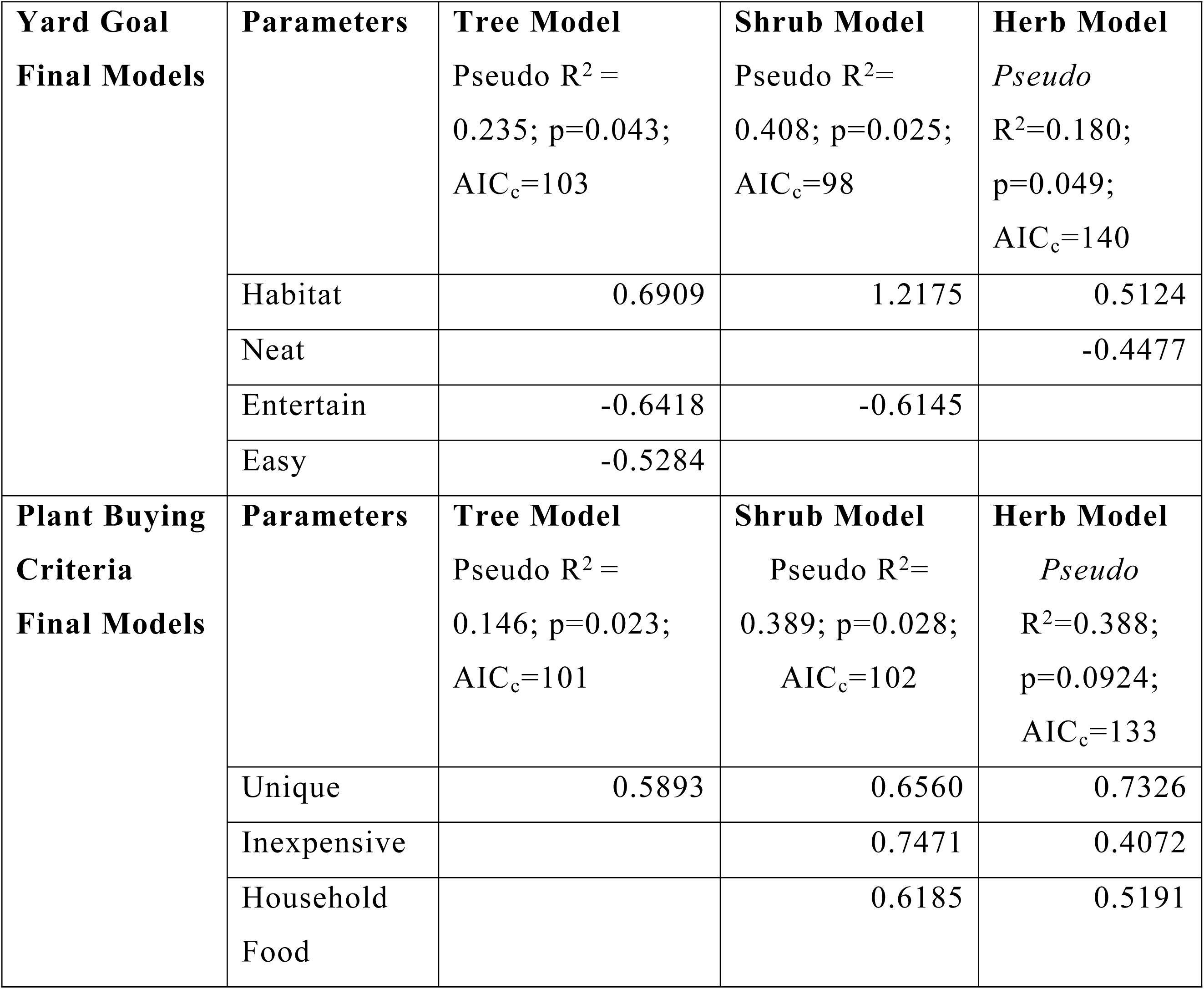
Standardized coefficients of homeowner planted species richness from best fitting negative regression models of homeowner preferences (n=34; S2). Corrected Akaike Information Criterion values (AIC_c_) within the same columns can be compared.

**Figure 6.**
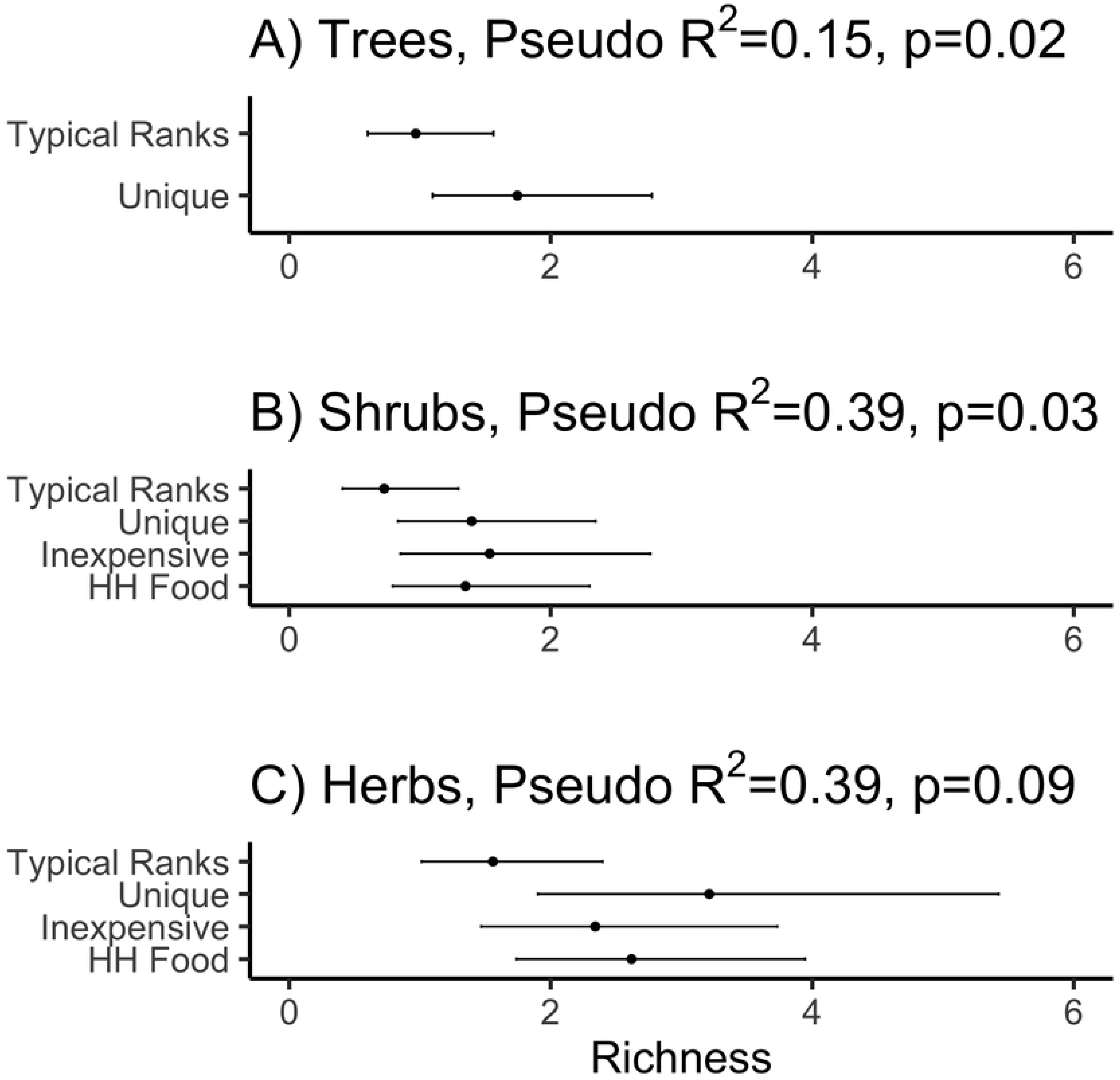
Estimated effects of higher rankings of plant selection criteria on homeowner planting behavior for A) trees, B) shrubs, and C) perennial herbs (n=34). Richness is the number of species that the homeowner planted. Closed circles represent the mean species richness resulting from homeowner plantings (Table 6; S3). Error bars are 95% confidence intervals. *Typical Ranks*: Typical homeowner ranking of criteria; *Unique* = Ranking Unique higher than typical; *Inexpensive* =Ranking Inexpensive higher than typical; *HH Food* = Ranking household food higher than typical.

**Figure 7.**
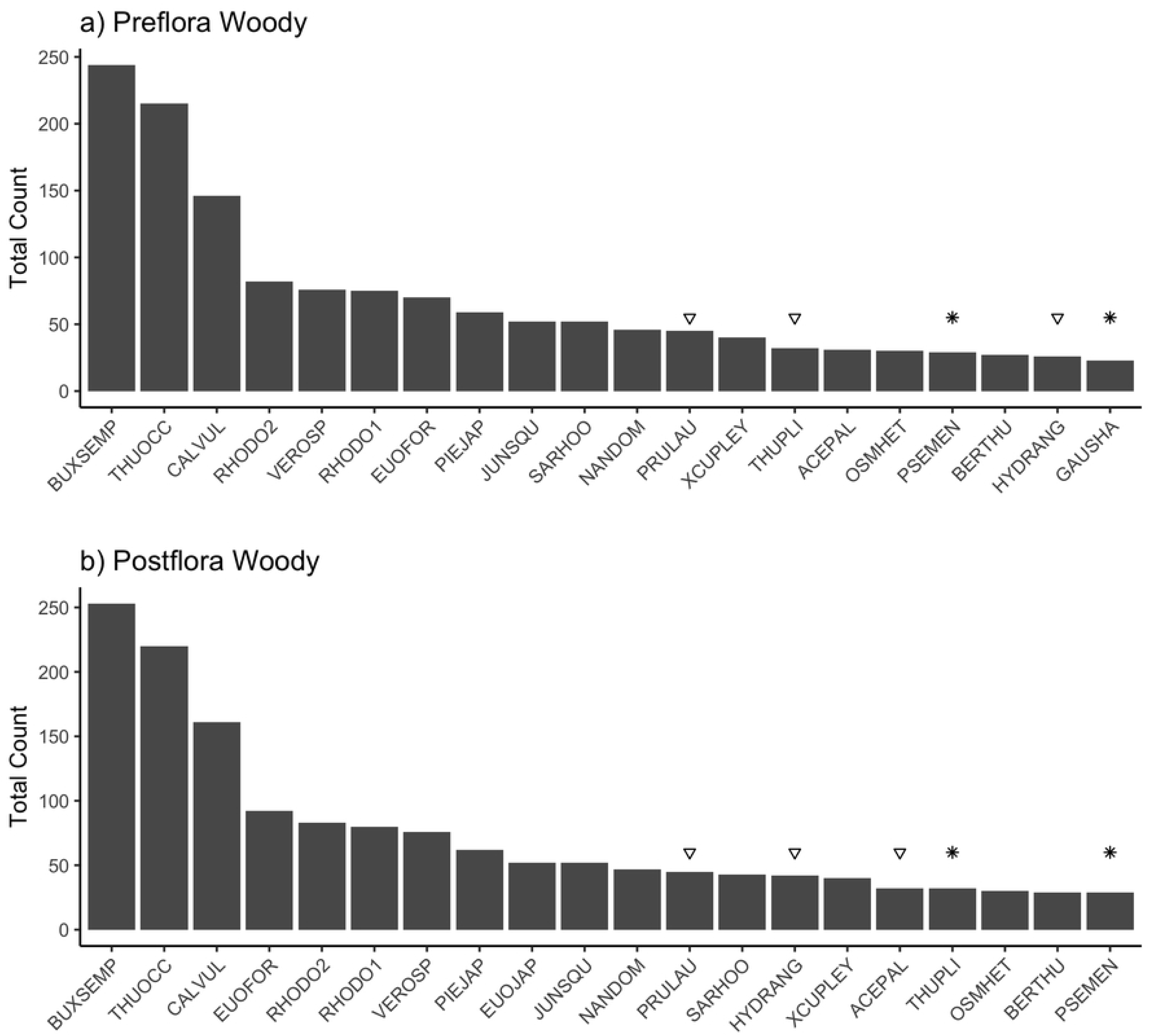
Counts of the 20 most abundant woody perennial taxa by flora. Species marked with an asterisk (*) are Pacific Northwest natives. Species marked with an upside triangle are deciduous trees or shrubs. All other species are evergreen trees or shrubs. ACEPAL= *Acer palmatum*, BERTHU=*Berberis thunbergii*, BUXSEM= *Buxus sempervirens*, CALVUL = *Calluna vulgaris*, EUOFOR=*Euonymus fortunei*, EUOJAP= *Euonymus japonicus*. GAUSHA = *Gaultheria shallon*, HYDRANG = *Hydrangea macrophylla*., JUNSQU = *Juncus squarrosa*, NANDOM = *Nandina domestica*, OSMHET= *Osmanthus heterophyllus*, PIEJAP= *Pieris japonica*, PRULAU = *Prunus laurocerasus*, PSEMEN=Pseudotsuga menziesii, RHODO1 = *Rhododendron* sp. 1, RHODO2=*Rhododendron* sp.2, SARCO=*Sarcococca* spp., THUOCC= *Thuja occidentalis*, THUPLI= *Thuja plicata*, VEROSP = *Veronica* sp., XCUPLEY = x*Cuprocyparis leylandii*.

Higher rankings of *Habitat* increased species richness for trees, shrubs, and herbs. In contrast, *Entertain* seemed to decrease species richness for trees and shrubs but not herbs. *Easy* decreased species richness for trees, and *Neat* decreased it for herbaceous perennials.

For trees, shrubs, and herbs, ranking *Unique* a higher than typical plant buying criterion increased species richness by about 1-3 species. Higher rankings of *Inexpensive* and *Household Food* also increased species richness for shrubs and for herbs.

### Homeowner plant removals

Through homeowner conversations, site evidence, and presale photos, I documented the removal or death of 39 trees, 22 shrubs, and 46 herbaceous perennials.

About half of the trees were evergreen and half were deciduous. Nearly 75% of the removed evergreen trees were improperly planted *Thuja occidentalis* (arborvitae) or *Thuja plicata* (western red-cedar), which were dead or dying when the homeowner moved in. Five were very large native evergreen trees (*P. menziesii* and *A. menziesii)*. Removals of living trees were to address specific homeowner problems and concerns. None were diseased or hazardous. One homeowner removed three large native trees, because they were “too close to the house.” Another reported removal of six developer-planted deciduous trees to fix a drainage problem in an otherwise inaccessible side yard. One homeowner removed a deciduous tree that was poorly sited, noting, “I got tired of whacking my head coming in from the mail.”

In contrast to removing living trees to fix specific site problems, homeowners reported removing other live perennials for other reasons: they did not like the plant, or they changed how the space is used. A homeowner replaced 12 graminoids in a front planting strip with short evergreen trees and herbaceous flowers, because “they were these really gross yellow grassy things.” Another removed five “weird, spiky plants.” Another homeowner reported that they “hate ivy” and removed it (*Hedera hibernica*) to plant a rock wall with “something more appropriate.” Two homeowners converted rain gardens to herb and vegetable beds.

In a particularly memorable case, a new homeowner removed front yard shrubs because she felt slighted by the builder. She complained that they gave her two very ugly shrubs “instead of a cool tree in the front yard.” By reviewing surrounding yards with her, I determined that the landscapers had planted two variegated *Euonymus fortunei* (winter creeper) instead of *Acer griseum* (paperbark maple). She reported that her house and yard were one of the last to be built in the large subdivision and that the model home landscaping was “way better” and had “lots more plants.” She plans to plant a tree to replace the ugly shrubs, but wants something “even cooler” than *A. griseum.* She said that she is still mad a year later.

### Changes in composition & abundance

Most species in the floras have fewer than 30 individuals (Figs 7,8). In general, woody perennial abundance changed less than that of herbaceous perennials..

**Figure 8.**
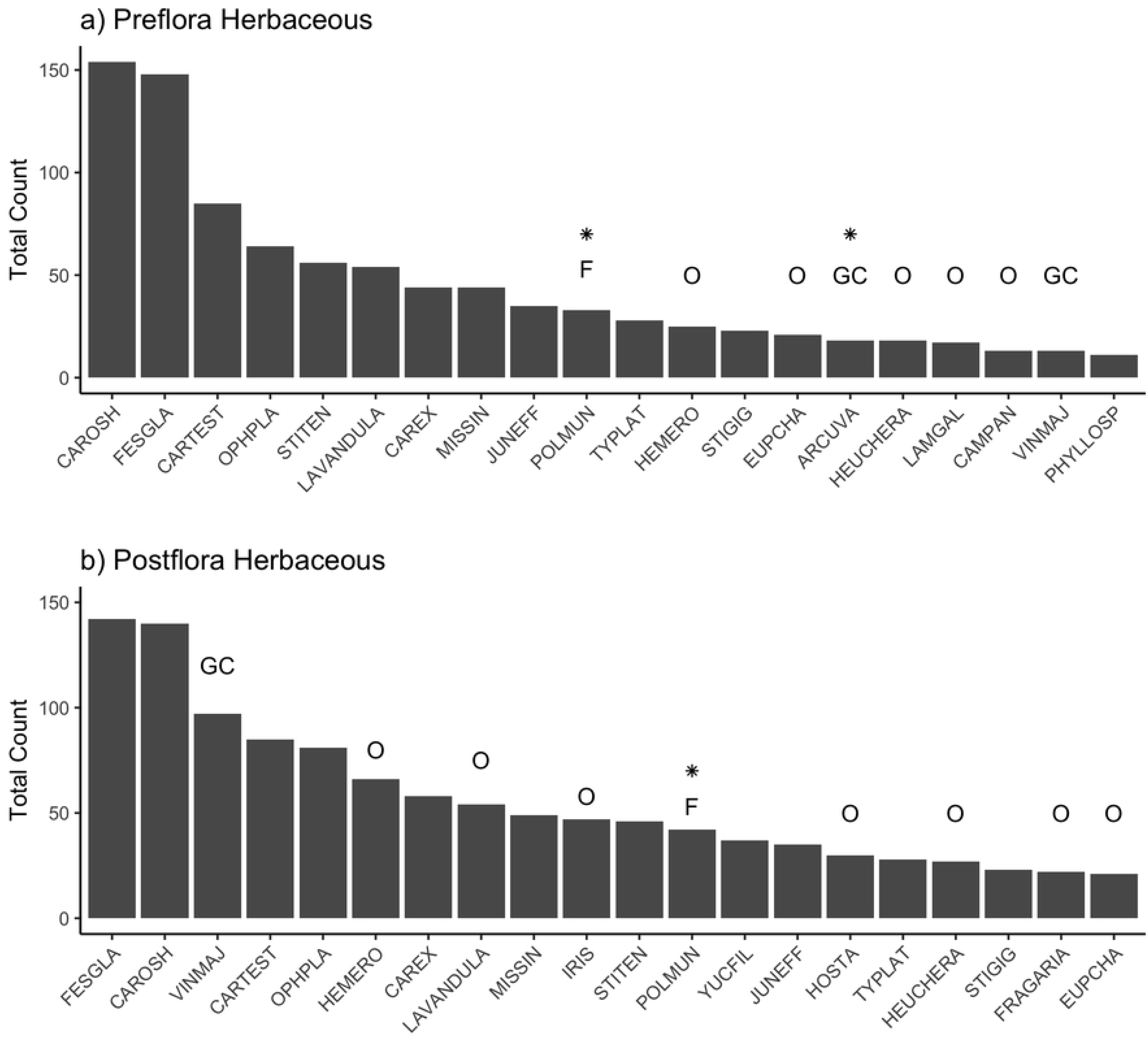
Counts of the 20 most abundant herbaceous perennial taxa by flora. Species marked with an asterisk (*) are Pacific Northwest natives. Unless marked by a species group (GC = groundcover; F = fern; O = other herbaceous perennial), herbs are graminoids. ARCUVA = *Arctostaphylos uva-ursi*, CAMPAN=*Campanula* spp., CAREX = *Carex elata* ‘Aurea’, CAROSH = *Carex oshimensis* ‘Evergold’, CARTEST=*Carex testacea*, EUPCHA= *Euphorbia characias*, FESGLA *= Festuca glauca*, FRAGARIA = *Fragaria* x *ananassa*, HEMERO = *Hemerocallis* spp., HOSTA = *Hosta* spp., HEUCHERA = *Heuchera* spp., IRIS = *Iris* spp., JUNEFF = *Juncus effusus*, LAVANDULA = *Lavandula* spp., LAMGAL= *Lamiastrum galeobdolon*, MISSIN = *Miscanthus sinensis*, OPHLA=*Ophiopogon planiscapus* ‘Nigrescens’, POLMUN = *Polystichum munitum*, STITEN = *Stipa tenuissima*, STIGIG = *Stipa gigantea*, TYPLAT = *Typha latifolia*, VINMIN = *Vinca minor*, YUCFIL = *Yucca filamentosa.*

*Buxus semperviren*s (common boxwood), *Thuja occidentalis (*arborvitae), and *Calluna vulgaris* (Scotch heather) are the most abundant woody perennials in both floras, ranging from about 150-250 individuals each. About 25% of the most abundant woody perennials are trees: *T. occidentalis*, x*Cuprocyparis leylandii* (Leyland cypress), *Acer palmatum* (Japanese maple), *Thuja plicata* (western red-cedar), and *Pseudotsuga menziesii* (Douglas-fir). Biggest changes in abundance of woody species are the appearance of *Euonymus japonicus* as the tenth most abundant woody perennial and the increase of *Hydrangea macrophylla* (bigleaf hydrangea) from 19^th^ to 14^th^ most abundant.

For herbaceous perennials, about half of the most abundant taxa are graminoids: *Carex oshimensis* (Oshima sedge), *Festuca glauca* (blue fescue), and *Carex testacea* (orange New Zealand sedge) have about 75-150 individuals each. *Lavandula* sp. (lavender) is the only dicotyledon in the top 10 most abundant herbs in the presale flora. In the postsale flora, four dicot herbs with showy flowers moved into the top 10: *Vinca minor* (periwinkle), *Hemerocallis* spp. (day lily), and *Iris* spp. (iris). In addition, *Hosta* spp. (hosta*), Yucca filamentosa* (Adam’s needle), and *Fragaria*.sp. (strawberry) appeared in the top 20 most abundant taxa.

The postsale flora is more taxonomically diverse than the presale flora, increasing from 131 species in 48 families to 200 species in 68 families (Table 7). Total herbaceous species richness increased the most (79%). Effective species richness (ESR) for all taxa ranged about 2-10 species in the presale flora and about 3-14 species in the postsale flora. Depending on perennial group (all, trees, shrubs, or herbs), this represents a 13-59% increase in alpha diversity. Beta diversity for all taxa is 10-19 presale and 111-22 postsale. Whether using species richness or Shannon to calculate beta from gamma diversity, beta diversity is largest for tree species in both floras.

**Table 7.**
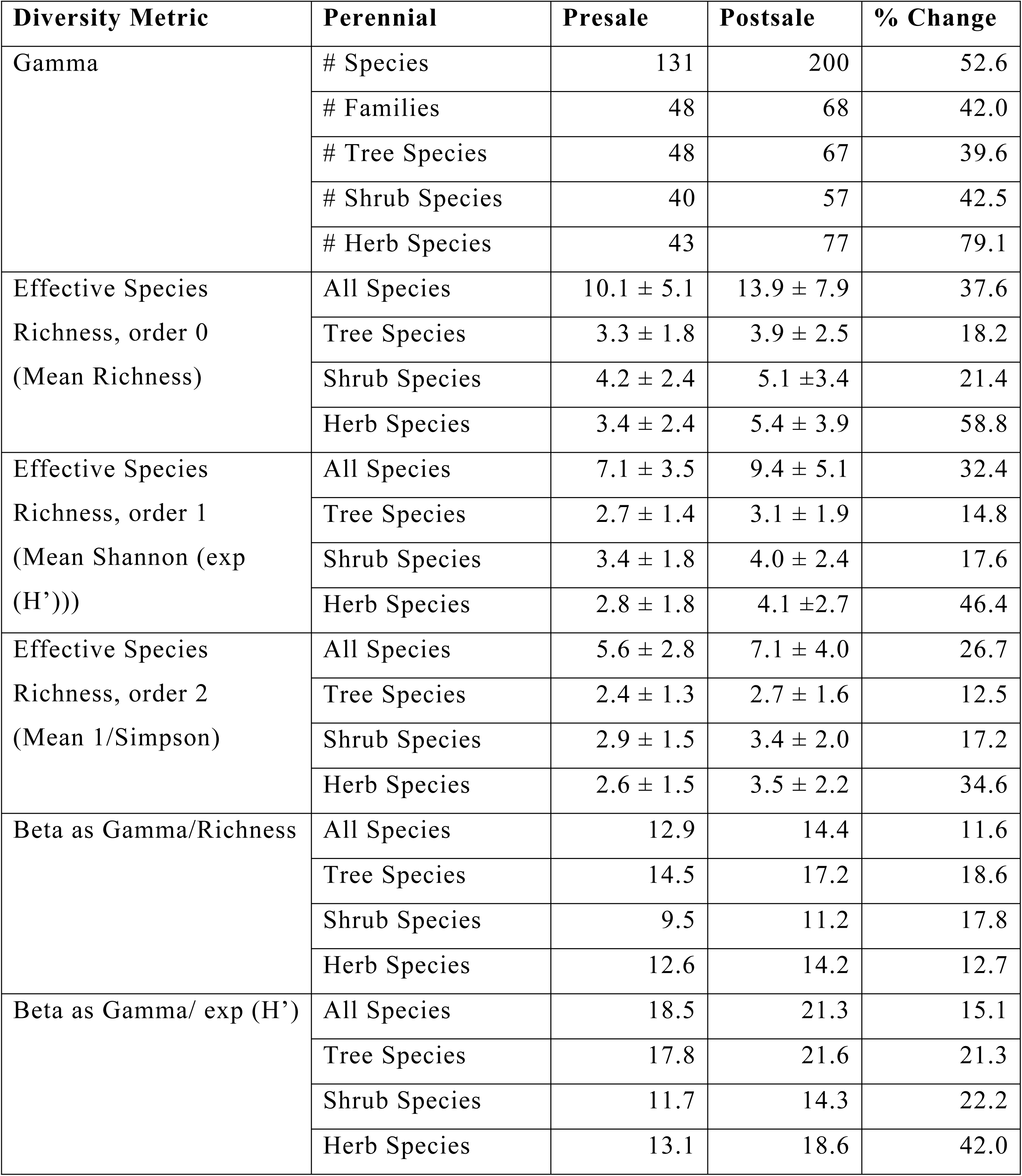
Perennial diversity by flora for all perennials and for trees, shrubs, and herbs (n=58). Errors are standard deviation.

| Diversity Metric | Perennial | Presale | Postsale | % Change |
| --- | --- | --- | --- | --- |
| Gamma | # Species | 131 | 200 | 52.6 |
|  | # Families | 48 | 68 | 42.0 |
|  | # Tree Species | 48 | 67 | 39.6 |
|  | # Shrub Species | 40 | 57 | 42.5 |
|  | # Herb Species | 43 | 77 | 79.1 |
| Effective Species<br>Richness, order 0<br>(Mean Richness) | All Species | 10.1 ± 5.1 | 13.9 ± 7.9 | 37.6 |
|  | Tree Species | 3.3 ± 1.8 | 3.9 ± 2.5 | 18.2 |
|  | Shrub Species | 4.2 ± 2.4 | 5.1 ± 3.4 | 21.4 |
|  | Herb Species | 3.4 ± 2.4 | 5.4 ± 3.9 | 58.8 |
| Effective Species<br>Richness, order 1<br>(Mean Shannon (exp<br>(H')))) | All Species | 7.1 ± 3.5 | 9.4 ± 5.1 | 32.4 |
|  | Tree Species | 2.7 ± 1.4 | 3.1 ± 1.9 | 14.8 |
|  | Shrub Species | 3.4 ± 1.8 | 4.0 ± 2.4 | 17.6 |
|  | Herb Species | 2.8 ± 1.8 | 4.1 ± 2.7 | 46.4 |
| Effective Species<br>Richness, order 2<br>(Mean 1/Simpson) | All Species | 5.6 ± 2.8 | 7.1 ± 4.0 | 26.7 |
|  | Tree Species | 2.4 ± 1.3 | 2.7 ± 1.6 | 12.5 |
|  | Shrub Species | 2.9 ± 1.5 | 3.4 ± 2.0 | 17.2 |
|  | Herb Species | 2.6 ± 1.5 | 3.5 ± 2.2 | 34.6 |
| Beta as Gamma/Richness | All Species | 12.9 | 14.4 | 11.6 |
|  | Tree Species | 14.5 | 17.2 | 18.6 |
|  | Shrub Species | 9.5 | 11.2 | 17.8 |
|  | Herb Species | 12.6 | 14.2 | 12.7 |
| Beta as Gamma/ exp (H') | All Species | 18.5 | 21.3 | 15.1 |
|  | Tree Species | 17.8 | 21.6 | 21.3 |
|  | Shrub Species | 11.7 | 14.3 | 22.2 |
|  | Herb Species | 13.1 | 18.6 | 42.0 |

## DISCUSSION

Residential perennials are a mix of native and non-native species that are either planted or else spontaneously established at different times. These floras can be viewed as assembling from a complex interaction of urban patterns and processes [7,8] or filters [9,40] that operate and interact at different scales. At the scale of a single parcel, perennials present in the yard represent the outcomes of decisions of local developers and homeowners about which species and individuals to keep, remove, or plant. Given that developers and homeowners have different incentives [8,85], understanding what each agent “typically” does and what each agent specifically did on a particular site is important to understand how these floras are assembled. Which plants do they keep from previous communities? Which species and types of plants do they choose to plant? How do those choices relate to the site and neighborhood context? Are homeowner yard management goals and plant selection criteria reflected in the species richness of their plantings?

### Characterizing developer decisions

Boone et al. [86] noted that vegetation in newly built suburbs is more likely to be the perception of what the developer believes will satisfy buyers, rather than an expression of the neighborhood or residents preferences. However, to subdivide land and to obtain building permits, residential developers must also comply with the local planning regulations [41,43,46]. Some of these directly affect the vegetation composition or structure. Some jurisdictions, but not all, have tree protection rules, specific tree species planting requirements, and incentives for builders to manage storm water on the parcel by creating rain gardens. Other planning regulations more indirectly influence vegetation. If a jurisdiction requires specific setback distance, then a developer now has a specific front yard area to landscape. Thus, new residential landscaping needs to be good enough for sale, satisfying both planners and potential buyers, but is less likely to reflect any particular household’s specific preferences.

To understand why residential developers choose “good enough” solutions, a phenomenon known as satisficing [87], Mohammed [85] incorporated lessons from behavioral economics [88-90], prospect theory [91,92], and mental accounting [93,94] to assess why developers might satisfice and focus on limiting costs. Broadly, given their risk aversion [95,96], relative changes in wealth are more influential in developer decision-making than absolute changes in wealth. Controlling costs after meeting predetermined project targets becomes more important than maximizing profit [97-99]. New residential landscaping, then, is more likely to be viewed as a cost to be controlled, rather than a way to increase sale price. Accordingly, many residential developers probably create the minimally acceptable landscape to meet planning regulations and expectations of target buyers.

The developer landscape satisficing decisions are a difficult to visualize legacy effect, but they lead to important outcomes in this study: 1) most of the native flora on a new parcel is from developer decisions, and 2) developers, not homeowners, are responsible for most of the woody perennials and graminoids. Because of developer preferences for evergreen trees, evergreen shrubs, and graminoids, along with a few reliable broadleaf species (e.g. *Acer* and *Lavandula* spp.), it is plausible that these perennials represent the safest, fastest way to create a finished, sellable landscape within the market in which they operate.

Other researchers have found neighborhood income to be positively associated with plant richness and abundance [10-12, 21-24]. In coining the luxury effect term, Hope et al. [10] noted, we don’t know whether “wealthier people create more diverse landscapes or simply acquire them.” Newly built and sold residential landscapes, therefore, offer a unique case to answer this question. Results from this study suggest that developers do target wealthier homebuyers by planting more diverse landscapes. Although appraised improvement value increased herbaceous perennial richness, parcel density more strongly increased herbaceous richness. If limited by planting area in more dense neighborhoods, developers may replace tree richness with herbaceous perennial richness.

However, developer behavior is heterogeneous, even accounting for planted area. The fact that the urban metric models explain <20% of the deviation suggests that developer planting decision heterogeneity needs further investigation.

### Characterizing homeowner decisions

Larsen and Harlan [49] draw on the work of Goffman [100], Nassauer [101], and Veblen [102] to describe residential landscapes as realized “presentations of social class.” Yards serve as transitional space, from public to personal, from the neighborhood to the household. Front yards represent the presented self, and backyards are the “personal pleasure ground.” Yards are therefore managed to meet the specific needs and interests of the particular household: spaces for playing, relaxing, entertaining, and gardening. After moving into their new home, the homeowners manage and change their yards and obtain plants in ways that depend on neighborhood norms and socioeconomic status [22, 103], household demographics [25,27,30,104-106], and how they intend to use the space [51,57,58].

Yet, most urban residential ecology research has attempted to link observed floral diversity patterns to household socioeconomics and urban gradients created from datasets at coarser scales than the parcel or household. Commonly used neighborhood surrogates to explore patterns in residential floral diversity or household management preference are census based boundaries [72,108,109] or Clarita’s, Inc’s. PRIZM system [105,106,110], which uses zipcode boundaries. I did detect a luxury effect for homeowner planted shrub richness, in the form of median assessed value, which explained about 7% of the deviation. However, plantable area, the strongest predictor of homeowner planted species richness for trees, shrubs, and herbs, still explained very little deviation. Individual household planting behaviors may not be easily predictable from coarsely aggregated socioeconomic measures. The multilevel models and the correlations suggest that household preferences and other factors more strongly influence landscaping behaviors than do urban drivers.

However, only a few studies have investigated the ways in which people’s preferences actually shape residential floral composition and diversity. In Australia, Kendal et al. [35] asked residents which traits they used to select plants and evaluated whether resident-expressed preferences for specific traits were reflected in their front yards. Specific plant-related traits included floral traits (flowering or not, flower size, color), leaf traits (size, shape, texture), plant form (shape, size, space filling). Resident preferences for these traits were evaluated with images of different plant species. They found that homeowner self-reported preferences for specific plant traits were weakly correlated with the actual plants in their front yards, that the strength of the correlations was stronger for residents who had lived in the home for at least five years, and that preferences for specific traits varied depending on which landscape type they had. Some traits used for plant selection were more related to plant management (low maintenance, drought tolerance, native) or specific useful functions (edibility, shade, screening, suppresses weeds, attracts birds, is fragrant), and others were more subjective (beauty, unusual, or personal reasons). Similarly, Cape Town, South Africa plant nursery shoppers reported selecting plants based on plant traits related to sensory appeal (appearance and scent), edibility, resource use (drought tolerance, ease of maintenance, native to area), and personal reasons relating to liking the plant, family history, or nostalgia [110]. In Salt Lake City, UT, residents named aesthetics, shade, ease of maintenance, size and shape, and fruit as top reasons that a particular tree in their yard was their favorite. *Acer* was the most frequently selected as a favorite genus [37].

In this study, homeowners’ preferences for using their new spaces were reflected in their planting decisions. Homeowners who ranked *Habitat* highly chose more diverse plantings, while ranking *Entertain, Easy*, or *Neat* highly led to less diverse plantings. Although *Habitat, Easy*, and *Neat* can be easily understood in terms of gardening interest, available resources, and concerns about yard appearance, *Entertain* is surprising. It could be that *Entertain* captures households who are satisfied with the overall structure of their yards and would rather use money and leisure time for socialization, rather than for gardening.

Homeowner plant buying criteria were also reflected in yards. *Unique* increased species richness for trees, shrubs, and herbs. *Household Food* and *Inexpensive* also increased species richness for shrubs and perennial herbs, but not trees. *Unique* and *Household Food* may be linked to gardening interest. *Inexpensive* suggests that homeowners who are price sensitive will buy and plant shrubs and herbs that are reduced in price.

Understanding reasons that homeowners remove plants requires thinking about ecosystem services and disservices [34], as well as considering how the unique household members want to use their new space or how they feel about particular plants. Removals may conflict with urban planning and conservation goals and incentives for preserving large urban trees [111,112] or managing urban stormwater via rain gardens [112-114]. Although developers left large trees on some parcels and constructed rain gardens on others, a few homeowners removed them soon after purchasing their homes.

The main reasons study participants removed perennials mostly related to perceived problems (death of improperly planted individuals, concerns about damage from really large trees, poorly sited trees), changing use of space, or really disliking certain species. Of these, only hazard concerns could be considered an ecosystem disservice. Although aesthetic traits are commonly cited as a reason to choose plants, it is problematic to link measures of beauty to observed patterns in residential landscapes, because of survivor bias. Very ugly or particularly hated plants are removed and are therefore never detected in preference studies.

These results are similar to other studies of tree removals. In Salt Lake City, UT [37], residents removed trees that died, were diseased, or were too big. A smaller proportion of homeowners removed trees because they didn’t like the particular species, or because they were poorly sited, was causing property damage, or required too much maintenance. Tenneson [53] found a similar list of reasons for tree removals in King County, WA: tree health and hazard concerns, property damage, maintenance needs, and size.

### Initial composition & abundance

I view these floras as an example of preemptive floristics [51,116]: whichever species and individuals are present in early stages tend to dominate, limiting the establishment and persistence of later arrivals. Therefore, whichever perennials the developer and homeowner establish initially sets the trajectory for other plant species might establish spontaneously as well as which plants someone else might plant later.

Evergreen perennials (Cupressaceae, Ericaceae, Pinaceae, Berberidacee, and Buxacaeae) and graminoids (Cyperaceae, Poaceae, Asparagaceae) dominate the floras. Other frequently represented families are Sapindaceae (*Acer* spp.), Rosaceae, and Lamiaceae. As the new homeowners began to manage their new space, they began to fill in the space according to their preferences and available resources. Thus, herbaceous perennials increased overall, as well as did the frequency of plant families with showy flowers: e.g. Rosaceae, Hydrangeaceae, Grossulariaceae, Iridaceae, and Asteraceae. Some of the frequency increase may be because the family is species rich (e.g. Rosaceae, Asteraceae), and some may be because new homeowners favor certain taxa (*Iris* spp. (Iridiaceae), *Hydrangea macrophylla* (Hydrangeaceae), *Ribes sanguineum* (Grossulariaceae). The J-shaped abundance curve I found for all is typical for residential floras, with most taxa occurring only rarely. For example, Jagamohan et al. [117] found that most species occurred in 5% or fewer of the 328 Indian yards they surveyed. They considered only six species to be common, occurring at more than 20% of sites. Similarly, Marco et al. [118] reported that 82% of the cultivated species in 120 gardens in southeastern France were represented by fewer than 50 individuals. In Trabazon, Turkey, Acar and Eroglu [119] found that 52/274 ornamental species occurred only once and that just 3 taxa occurred at 45% or more sites: *Rosa* sp., *H. macrophylla*, and *Nerium oleander*.

Other studies of residential landscapes found more diversity. In 163 yards in Los Angeles, CA, USA, Clarke et al. [11] found 109 tree species, with alpha diversity (exp H’) of 4.17. In Salt Lake City, UT, USA, Avolio et al. [36] found 132 yard tree species; mean species richness was 3.4 ± 0.4 for low income yards and 6.8±0.6 for higher income yards. In Catalonia, Spain, Cubino et al. [119] found 630 plants from 245 yards; mean species richness was about 35 species/yard. In Turkey, Acar and Eroglu [120] reported mean species richness of 8.59 ±3.76 for traditional housing and 33.37±23.43 for public housing for employees. However, these studies represent a mix of ages and are not easily comparable the very new yards in this study.

Although vegetation cover in my sample will likely increase over time from vegetative growth, it is unclear how much species diversity may change. Given that most of the increase in species richness from homeowner plantings is from herbaceous perennials, it seems likely that species richness of woody perennials would stay relatively stable but that species richness of herbaceous perennials may continue to increase. Summit and McPherson [28] found that homeowners planted the most trees within the first 5 years of tenure. Planting rates for shrubs and herbaceous perennials are not well documented in the literature and need further scrutiny.

Urban species pools in general and residential landscapes in particular support a large proportion of non-native species (9,121]. In this study, all three species pools (remnant, developer, and homeowner) consist mostly of non-native perennials. The proportion of non-native taxa for the remnant (61%) and developer (79%) pools is within the range found for other studies of residential landscapes, but the household selected proportion (90%) is greater than in other studies. For European yards, non-native taxa comprised 70% of the flora in United Kingdom studies [122,123], 80% in southeastern France [112], and 68-77% in Spain [124,125]. Based on surveys of Puerto Rican yards, residents, and nurseries, Torres-Camacho et al. [106] found that a multitude of interacting factors shape the relative frequency of native plants in residential landscapes: the nursery industry having few available natives, historical plantings, informal exchanges of plants from friends and family, and natural dispersion processes.

### Strengths & limitations

Although the small size limits generalizability, the sample is a drawn from a larger probability sample of newly built homes and yards. This cohort-based approach allowed investigation of the different choices that people make within the same real estate market. I chose to deliberately oversample parcels developed by small firms, in order to evaluate a variety of decisions of development firms and homeowners who choose different locations and development types across the urban gradient. Homeowners who are more interested in gardening may have been more likely to give permission, so results could be biased towards those homeowners who have an interest in plants.

The multilevel models validated the use of using plant characteristics linked to how plants are actually recommended and sold to consumers in large retail stores and nurseries. Some of these plant structure groups are commonly used in the residential landscape literature (e.g. leaf habit, height), while others relating to herbaceous perennials are not. By accounting for missing structure groups in planting decisions, I found distinct differences in developer and homeowner behaviors and preferences for certain groups overall. Because developers planted most of the woody vegetation, other urban ecology researchers should consider how they separate homeowner retention decisions from planting decisions. Otherwise, researchers may attribute urban ecosystem drivers of woody perennial composition and diversity to the wrong agent.

Because the urban drivers at the parcel and neighborhood extents explained <20% of deviations, it seems likely that individual preferences and other constraints of developers and homeowners also drive species richness patterns. Because I had homeowner preference data, I was able to find stronger predictors of household planted species richness. However, I did not have similar preference data for developers.

Removals are the most error prone of the three actions (retention, planting, removal), because they are not easy to detect. The true number of removals is likely to be larger. Plant identifications of removed plants were much better for woody perennials. Stumps were easy to see, and larger plants were more obvious in photos. Removal of woody species also seemed more memorable to homeowners, because of the effort, time, and money involved.

## CONCLUSIONS

This study is one of the first assessments of how developer and homeowner decisions together shape the flora of residential landscapes. Using plant structural groups to reflect ways that plants are listed in gardening literature and marketed in stores and nurseries allowed me to detect distinct differences in the types of plants each agent chose. More research on new residential landscapes in other regions is needed to assess variability of landscaping decisions of the developers and homeowners.

Plant choice heterogeneity has important implications for urban planners, ecologists, and modelers, and others who seek to understand urban landscape change. First, developers are responsible for most of the woody perennials and graminoids, and homeowners are responsible for more of the other herbaceous perennials. Researchers should ask current homeowners or residents to identify which perennials they have planted and which were already present. Second, most of the native flora results from developer decisions. Conservation efforts to increase native flora on private lands may require work with stores, nurseries, and households in specific locations to increase native flora availability and desirability. Third, planning regulations and incentives in some jurisdictions led to developers retaining large trees and installing rain gardens to manage storm water on site. However, this does not mean that the new homeowners keep the large trees or the rain gardens. Urban greening goals and strategies may need to be adjusted to account for potential removals. Fourth, longer term cohort studies are needed to understand vegetation dynamics across urban landscapes and within cohort heterogeneity.

A large body of research links neighborhood wealth data to patterns of plant species richness, particularly for woody perennials. Most homeowners did not plant trees and shrubs; developers did. Therefore, it is the retention decisions of homeowners, rather than their planting decisions, driving a large proportion of woody species richness in yards. Landscaping decisions appear to be driven by how homeowners want to use their yards and how they select plants. More work is needed to understand planting decisions of developers.

## Acknowledgements

I thank the 60 participating homeowners who granted written permission to sample their yards and told me what they changed. Kimberly Trask, Jonnie Narita, Sophia O’Hara, Elena Boyle, and Ryan Boyle helped measure area and count plants. Richard Olmstead reviewed nomenclature and confirmed unusual plant identifications. Marina Alberti, Jan Whittingon, David Montgomery, and members of the Urban Ecology Research Lab provided additional scientific guidance. Yicheng Li from the University of Washington Center for Statistics and the Social Sciences provided statistical consulting services.

## Supporting information

**S1. Perennial names, structure group classification, and frequencies.** CODE: Species code; Name: Scientific name; Genus: Plant Genus, Family: Plant Family, PNW: 1 = native to Pacific Northwest and 0 = non-native; Grp: Plant Structure Group [51]. dev = # of sites that developer planted perennial species; hh: # of sites that homeowner planted species; remnant: # of sites remnant perennial occurred.

**S2. Supplementary data tables for multilevel models (S2 Tables 1,2), urban metrics (S2 Tables 3-8), and household preferences (S2 Tables 9-12).**

